# Plasma β-hydroxybutyrate Concentrations in Young Adult Females After a High-Fat Meal Under Normoxemia, Intermittent Hypoxemia, and Continuous Hypoxemia

**DOI:** 10.64898/2026.03.11.711039

**Authors:** Nicholas Goulet, Alexanne Larocque, Caroline Marcoux, Vincent Bourgon, Jean-François Mauger, Ruwan Amaratunga, Pascal Imbeault

## Abstract

Hypoxemia occurs in intermittent forms, such as obstructive sleep apnea, and in continuous forms, such as at high altitude, and is increasingly recognized as a modulator of cardiometabolic risk. Although hypoxemia alters postprandial glucose and lipid metabolism, its effects on ketone bodies remain unclear. Using a randomized crossover design, we examined whether six hours of normoxemia or intermittent hypoxemia (15 hypoxemic cycles/hour targeting ∼85% peripheral oxyhemoglobin saturation with 100% medical-grade nitrogen) alters plasma β-hydroxybutyrate (BHB) concentrations in 12 young adult females (mean [SD]: 21 [3] years) following a high-fat meal (33% of estimated daily energy requirements; 59% of calories from fat). In a follow-up session, a subset (n = 8) completed six hours of continuous hypoxemia (fraction of inspired oxygen ∼12.0% in a normobaric chamber). Postprandial data were analyzed using baseline-adjusted linear mixed-effects models, with Bonferroni post hoc tests. A time × condition interaction (P = 0.010) indicated that BHB concentrations at 360 minutes were higher during continuous hypoxemia (0.247 mmol/L; 95% CI: 0.218-0.275) than normoxemia (0.176 mmol/L; 95% CI: 0.153-0.200; P_Bonferroni_ = 0.029) and intermittent hypoxemia (0.163 mmol/L; 95% CI: 0.139-0.186; P_Bonferroni_ = 0.002), representing increases of 13.0% and 14.2% in estimated marginal means, respectively. This response was accompanied by higher postprandial plasma glucose and triglyceride concentrations during continuous hypoxemia than during normoxemia and intermittent hypoxemia (P_Bonferroni_ ≤ 0.002), despite similar plasma insulin and non-esterified fatty acid responses across conditions (P ≥ 0.081). These findings indicate that continuous hypoxemia increases late postprandial plasma BHB concentrations in young adult females.

**New Findings:** *What is the central question of this study?:* What are the effects of normoxemia, intermittent hypoxemia, and continuous hypoxemia on plasma β-hydroxybutyrate (BHB) concentrations in young adult females after a high-fat meal?

*What is the main finding and its importance?:* Compared to normoxemia, young adult females showed higher postprandial plasma BHB concentrations during continuous hypoxemia, but not during intermittent hypoxemia, despite similar changes in plasma concentrations of two main regulators of BHB production (non-esterified fatty acids and insulin) across experimental conditions. These findings suggest that continuous hypoxemia modifies postprandial BHB concentrations through mechanisms not fully explained by circulating non-esterified fatty acids or insulin concentrations alone.

## 1. Introduction

Hypoxemic conditions, such as obstructive sleep apnea and high altitude, are increasingly recognized as modifiers of cardiometabolic risk, in part through their effects on postprandial metabolism (Chopra et al., 2017; Woolcott et al., 2015). Recurrent or sustained reductions in arterial oxygen saturation may alter substrate utilization, insulin sensitivity, and lipid metabolism, processes that are particularly relevant during the postprandial period (Morin et al., 2021). While hypoxemia-related alterations in postprandial glucose and triglyceride metabolism are being increasingly described, their effects on postprandial ketone body dynamics in humans remain relatively unexplored. This represents an important knowledge gap, as ketone bodies may partly serve as signalling molecules that modulate cardiometabolic risk under oxidative or hypoxic stress (Stalmans et al., 2024).

β-hydroxybutyrate (BHB) is the predominant circulating ketone body and provides an index of hepatic lipid-derived substrate handling (Newman & Verdin, 2017). Because BHB reflects the balance between hepatic fatty acid delivery, insulin-mediated suppression of ketogenesis, and mitochondrial oxidative capacity, it may capture aspects of metabolic regulation that are not apparent from glucose or triglyceride responses alone (Cotter et al., 2013; Puchalska & Crawford, 2017; Veech, 2004). Laboratory-based experimental models of continuous hypoxemia (e.g., simulated high-altitude) and intermittent hypoxemia (e.g., simulated obstructive sleep apnea) have demonstrated measurable effects on postprandial glucose and lipid metabolism (Goulet, Marcoux, et al., 2024; Goulet, Morin, et al., 2024; Morin et al., 2021). However, studies examining postprandial BHB responses to hypoxemia are sparse, especially in humans, and direct comparisons between intermittent and continuous hypoxemic exposures are lacking.

Our group previously demonstrated that, compared to normoxemia, continuous hypoxemia increased fasting plasma BHB concentrations over six hours; however, it did not elevate BHB concentrations during a constant feeding period (Marcoux et al., 2022). Additionally, BHB concentrations remained statistically similar after six hours of intermittent hypoxemia under a postprandial state, compared with normoxemia (Marcoux et al., 2022). Notably, these findings were limited to young adult males and did not directly compare intermittent and continuous hypoxemic exposures under matched nutritional states. In parallel, we have demonstrated that females exhibit attenuated postprandial disturbances in glucose and lipid metabolism in response to intermittent hypoxemia (Goulet, Marcoux, et al., 2024). Building on this prior work, the present study examined plasma BHB responses, along with plasma concentrations of other metabolites (glucose, non-esterified fatty acids [NEFA], triglycerides) and insulin, during six hours of normoxemia, intermittent hypoxemia, and continuous hypoxemia in young adult females following the consumption of a high-fat meal. We considered this an exploratory study to generate hypotheses for future research. As such, there were no formal hypotheses.

## 2. Methods

### 2.1. Ethical approval

This single-site, laboratory-based, randomized crossover study was approved by the University of Ottawa Health Sciences and Science Research Ethics Board (ethics file number: H-06-18-837) and conducted in accordance with the Declaration of Helsinki, except for prospective registration in a publicly accessible database. All participants provided written informed consent before enrollment. The present study was conducted as part of a larger randomized crossover trial examining the effects of hypoxemia on postprandial metabolism in humans; detailed descriptions of the preliminary screening procedures and sample size considerations have been published previously (Goulet, Marcoux, et al., 2024); key methodological elements relevant to the current analyses are reiterated here for completeness.

### 2.2. Participants

Participants were recruited between April 2022 and April 2023 using convenience sampling at the University of Ottawa (Ottawa, Canada). Twelve young adult females met the eligibility criteria and completed the experimental protocol. Inclusion criteria were age 18-30 years and habitual sleep duration of 7-10 hours per night. Exclusion criteria included a history of respiratory disease, hypertension, cardiovascular disease, diabetes, current tobacco smoking, use of lipid-lowering medication, use of hormonal contraceptives, pregnancy, lactation, irregular menstrual cycles, and lactose intolerance (which would have precluded consumption of the high-fat meal).

### 2.3. Experimental design

Each participant completed one screening session (described in section 2.4) followed by two experimental sessions conducted under normoxemia and intermittent hypoxemia in a randomized crossover order (described in section 2.5). A third experimental session involving continuous hypoxemia was subsequently completed by a subset of participants as part of a follow-up session (described in section 2.6). Participants were instructed to obtain at least seven hours of sleep and to abstain from exercise, caffeine, alcohol, and anti-inflammatory medications for 36 hours before each experimental session. A standardized dinner was provided and consumed approximately 12 hours before arrival at the laboratory. The meal consisted of meat lasagna or vegetarian macaroni and cheese (President’s Choice, Canada), with a gluten-free alternative provided to one participant with celiac disease. Nutritional composition was comparable across meals (Goulet, Marcoux, et al., 2024). Experimental sessions began between 06:00 and 09:00 (consistent within-participant), conducted during the early follicular phase of the menstrual cycle (one to seven days following self-reported menstruation), and were separated by at least one menstrual cycle. Female steroid sex hormones were measured to confirm that serum concentrations were similar between the experimental conditions and within the normal ranges for the early follicular phase (Stricker et al., 2006).

### 2.4. Screening session

During the screening session, participants completed medical and lifestyle questionnaires, including the Pittsburgh Sleep Quality Index. Height and body weight were measured using a stadiometer (Perspective Enterprises, USA) and a beam scale (Tanita, USA), respectively. Body composition was assessed using dual-energy X-ray absorptiometry (GE Lunar Prodigy, USA). Basal metabolic rate was measured by indirect calorimetry (VIASYS Healthcare Inc., USA) following a 12-hour overnight fast in a thermoneutral, darkened environment, with participants resting supine for 30 minutes. Daily energy expenditure was estimated using the Harris–Benedict equation by multiplying the measured basal metabolic rate by a physical activity factor of 1.375. Participants were also exposed to intermittent hypoxemia (≤ 20 minutes) using the experimental apparatus described below and monitored for symptoms of acute mountain sickness using the Lake Louise scoring system. All participants tolerated the exposure without adverse events.

### 2.5. Experimental sessions: normoxemia and intermittent hypoxemia

Upon arrival for each experimental session, resting blood pressure was measured using an automated sphygmomanometer (American Diagnostic Corporation, USA). An intravenous catheter was inserted into an antecubital vein, and a baseline blood sample was collected. Participants then consumed a high-fat meal providing 33% of estimated daily energy expenditure (59% fat, 31% carbohydrate, 10% protein), composed of Ensure Plus (Abbott Laboratories, USA) and 35% whipping cream (Sealtest, Canada). Water was available ad libitum throughout the session. Participants remained awake in a semi-recumbent position for six hours and were continuously monitored using a fingertip pulse oximeter (Masimo, USA), with heart rate and peripheral oxygen saturation (SpO_2_) recorded at 1-second intervals.

Following meal consumption, participants were exposed for six hours to normoxemia (ambient air: fraction of inspired oxygen [FiO_2_] ∼20.93%, SpO_2_ ∼98%) or intermittent hypoxemia, consisting of 15 hypoxemic cycles/hour, achieved by inhalation of 100% medical-grade nitrogen via an oronasal mask with a two-way non-rebreathing valve. Nitrogen inhalation continued until SpO_2_ reached 85%, after which the inspiratory line was switched to ambient air, where SpO_2_ returned to baseline before initiating the next cycle. This frequency approximates a moderate apnea-hypopnea index (15 events/hour), a common metric of obstructive sleep apnea severity. The order of experimental conditions was randomized, with four participants completing the intermittent hypoxemia condition first. Mild symptoms of acute mountain sickness were reported by one participant following intermittent hypoxemia and by one participant following normoxemia.

### 2.6. Follow-up session: continuous hypoxemia

After completing the primary trial, participants were invited to complete an additional experimental session involving continuous hypoxemia. Eight participants consented to this follow-up; four declined due to scheduling constraints. All pre-experimental procedures were identical to the primary protocol, except that sessions were not restricted to a specific menstrual cycle phase. During this condition, participants were exposed for six hours to continuous normobaric hypoxia simulating ∼5000 m altitude (FiO_2_ ∼12.0%; SpO_2_ ∼80%) in a climate-controlled hypoxic chamber with oxygen extractors (Altitude Control Technologies, USA). Two participants reported mild symptoms consistent with acute mountain sickness during this exposure.

### 2.7. Blood sample collection

Venous blood samples were collected at baseline and 30, 60, 90, 120, 180, 240, 300, and 360 minutes post-meal into EDTA-treated and serum tubes (BD Vacutainer, USA). Plasma and serum were separated by centrifugation at 1,250 RCF for 10 minutes at 4°C. Serum samples were allowed to clot for ∼30 minutes at room temperature before centrifugation. Aliquots were stored at -80°C until analysis.

### 2.8. Analyte assays

Serum concentrations of progesterone and β-estradiol were measured at baseline, whereas plasma concentrations of glucose, NEFA, triglycerides, and insulin were measured at baseline and 30, 60, 90, 120, 180, 240, 300, and 360 minutes post-meal. Plasma BHB concentrations were measured hourly (baseline and 60-360 minutes post-meal) and were not assessed at the 30- or 90-minute time points. All samples were quantified in duplicate using commercially available colorimetric assays, following the manufacturer’s protocols, as previously described by our group (Goulet, Marcoux, et al., 2024; Marcoux et al., 2022).

### 2.9. Statistical analysis

Participant characteristics between the primary group and the follow-up subgroup were compared using unpaired t-tests. Physiological variables (blood pressure, heart rate, and SpO₂), serum concentrations of progesterone and β-estradiol, and baseline plasma concentrations of BHB and other metabolites (glucose, NEFA, and triglycerides) and insulin were analyzed using linear mixed-effects models with condition (normoxemia, intermittent hypoxemia, and continuous hypoxemia) as a fixed effect and participant identification as a random intercept. Postprandial concentrations of plasma BHB, other metabolites, and insulin were analyzed using linear mixed-effects models with condition and time (30, 60, 90, 120, 180, 240, 300 and 360 minutes post-meal) as fixed effects, and participant identification as a random intercept, while baseline (0 minutes) was included as a continuous covariate. Differences in BHB concentrations between baseline and 360 minutes post-meal were analyzed using a linear mixed-effects model with time and condition as fixed effects and participant identification as a random intercept.

Descriptive data for each condition are presented as means and standard deviations of raw data. Within- and between-condition differences are presented as model-derived estimated marginal means with 95% confidence intervals (CI). Bonferroni’s post hoc comparisons were performed when a significant main effect or interaction was identified. Homoskedasticity and normality of the residuals were assessed using diagnostic plots. Statistical analyses were performed in *jamovi* (v2.5.6; *gamlj3* module v3.3.4), with an alpha level of 0.050. Figures were generated using GraphPad Prism (v10.6.1).

## 3. Results

Participant characteristics at enrollment for the primary group and the follow-up subgroup are presented in Table 1. No differences were observed in any characteristic between groups (all P ≥ 0.532). Baseline serum concentrations of progesterone and β-estradiol are presented in Table 2 and were similar across experimental conditions (both P ≥ 0.155).

**Table 1.**
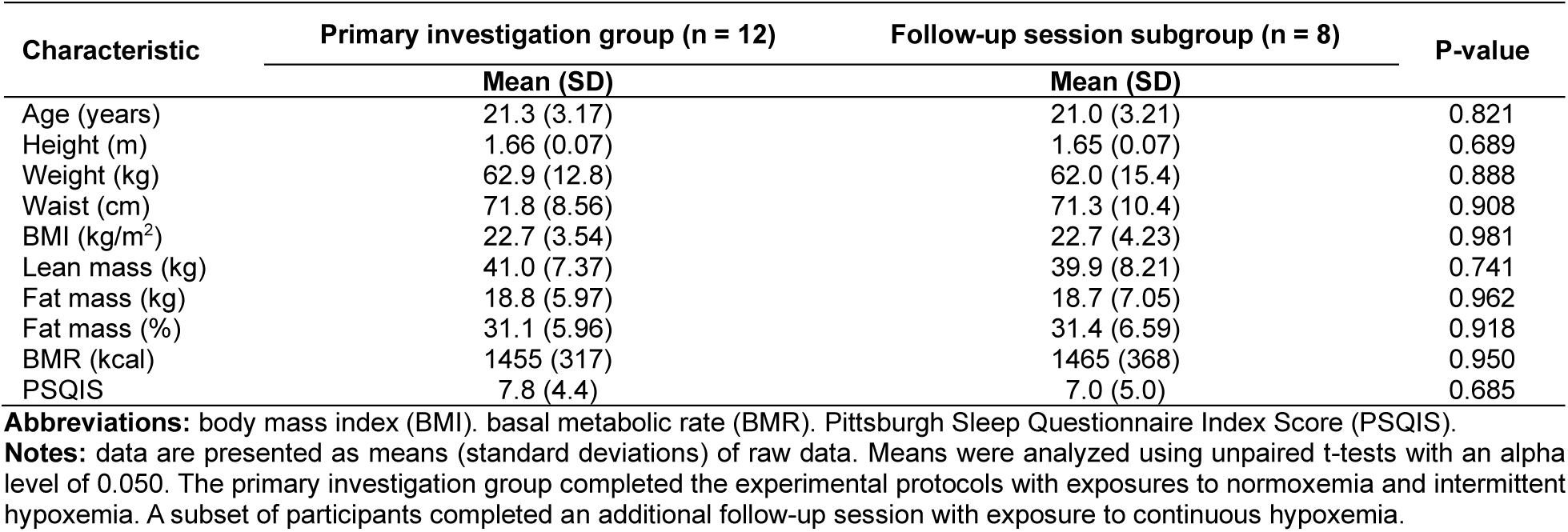
Characteristics of the primary investigation group and the follow-up session subgroup.

| Characteristic | Primary investigation group (n = 12) | Follow-up session subgroup (n = 8) | P-value |
| --- | --- | --- | --- |
|  | Mean (SD) | Mean (SD) |  |
| Age (years) | 21.3 (3.17) | 21.0 (3.21) | 0.821 |
| Height (m) | 1.66 (0.07) | 1.65 (0.07) | 0.689 |
| Weight (kg) | 62.9 (12.8) | 62.0 (15.4) | 0.888 |
| Waist (cm) | 71.8 (8.56) | 71.3 (10.4) | 0.908 |
| BMI (kg/m<sup>2</sup>) | 22.7 (3.54) | 22.7 (4.23) | 0.981 |
| Lean mass (kg) | 41.0 (7.37) | 39.9 (8.21) | 0.741 |
| Fat mass (kg) | 18.8 (5.97) | 18.7 (7.05) | 0.962 |
| Fat mass (%) | 31.1 (5.96) | 31.4 (6.59) | 0.918 |
| BMR (kcal) | 1455 (317) | 1465 (368) | 0.950 |
| PSQIS | 7.8 (4.4) | 7.0 (5.0) | 0.685 |

**Abbreviations:** body mass index (BMI), basal metabolic rate (BMR), Pittsburgh Sleep Questionnaire Index Score (PSQIS).
**Notes:** data are presented as means (standard deviations) of raw data. Means were analyzed using unpaired t-tests with an alpha level of 0.050. The primary investigation group completed the experimental protocols with exposures to normoxemia and intermittent hypoxemia. A subset of participants completed an additional follow-up session with exposure to continuous hypoxemia.

**Table 2.** Serum concentrations of progesterone and β-estradiol per experimental condition.

| Steroid sex hormone | Normoxemia | Intermittent hypoxemia | Continuous Hypoxemia | P-value |
| --- | --- | --- | --- | --- |
| Progesterone (ng/mL) | 1.65 (1.6) | 2.76 (4.2) | 5.00 (5.2) | 0.155 |
| Estradiol (pg/mL) | 90.7 (47) | 92.2 (42) | 73.5 (47) | 0.244 |

**Notes:** data are presented as means (standard deviations) of raw data and were compared using linear mixed-effects models with condition as a fixed effect and participant identification as a random intercept with an alpha level of 0.050. The analyses of serum β-estradiol concentrations exclude one outlier (normoxemia: 652 pg/mL; intermittent hypoxemia: 822 pg/mL), who had values > 14 SD above the group mean. The same participant also displayed serum progesterone concentrations above the 95th percentile (normoxemia, 5.41 ng/mL; intermittent hypoxemia, 6.00 ng/mL) (Stricker et al., 2006).

### 3.1. Physiological responses to six hours of exposure to normoxemia, intermittent hypoxemia, and continuous hypoxemia

Table 3 presents the mean systolic and diastolic blood pressures, heart rate, and SpO_2_ during normoxemia, intermittent hypoxemia, and continuous hypoxemia. Systolic and diastolic blood pressures were similar between normoxemia, intermittent hypoxemia, and continuous hypoxemia (both P ≥ 0.201). Heart rate differed across conditions (P < 0.001), with higher values observed during continuous hypoxemia (95.0 bpm; 95% CI: 87.5-102.5) compared with normoxemia (76.9 bpm; 95% CI: 70.6-83.2; P_Bonferroni_ < 0.001) and intermittent hypoxemia (79.8 bpm; 95% CI: 73.5-86.1; P_Bonferroni_ < 0.001), whereas normoxemia and intermittent hypoxemia did not differ (P_Bonferroni_ = 0.631). Mean SpO_2_ also differed across experimental conditions (P < 0.001), being lowest during continuous hypoxemia (83.9%; 95% CI: 82.2-85.6) compared with normoxemia (97.8%; 95% CI: 96.7-98.9; P_Bonferroni_ < 0.001) and intermittent hypoxemia (96.1%; 95% CI: 94.9-97.2; P_Bonferroni_ < 0.001), which did not differ from each other (P_Bonferroni_ = 0.102). Time spent below SpO_2_ thresholds (≤ 90%, ≤ 85%, and ≤ 80%) increased stepwise from normoxemia to intermittent hypoxemia to continuous hypoxemia (all P < 0.001).

**Table 3.**
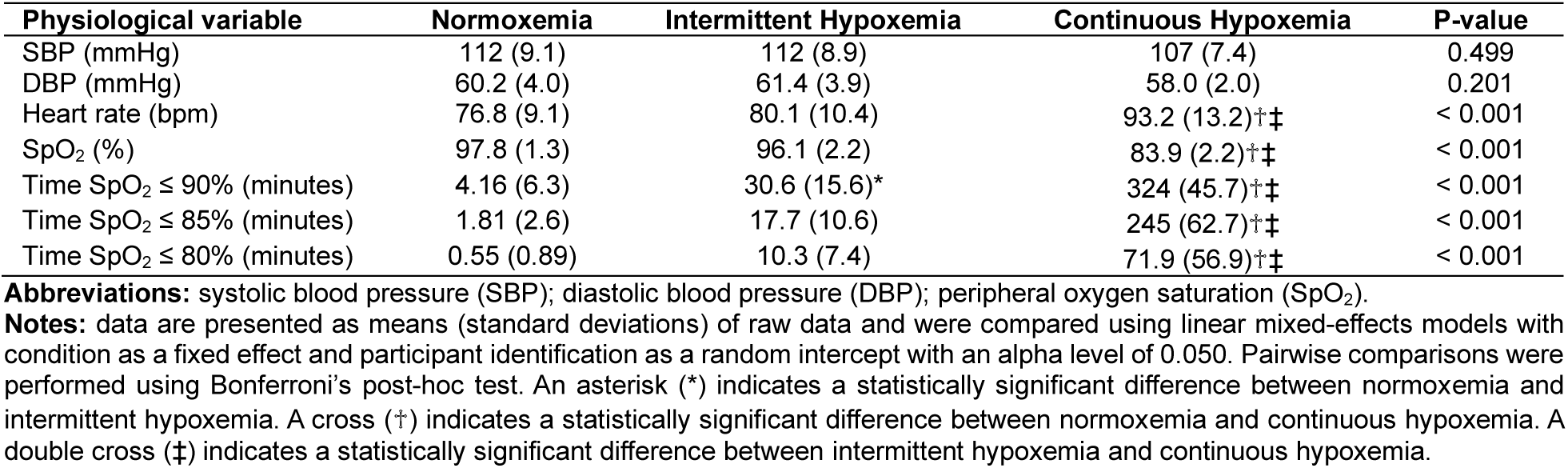
Mean systolic and diastolic blood pressures, heart rate, and peripheral oxygen saturation following the consumption of a high-fat meal during six hours of exposure to normoxemia, intermittent hypoxemia, and continuous hypoxemia in young adult females.

### 3.2. Plasma BHB response to a high-fat meal during six hours of exposure to normoxemia, intermittent hypoxemia, and continuous hypoxemia

Figure 1A presents baseline and postprandial plasma BHB concentrations. Baseline plasma BHB concentrations were similar across experimental conditions (P = 0.167). A time × condition interaction was observed for postprandial BHB concentrations (P = 0.010), indicating that the temporal pattern of the BHB response differed across conditions. Within-condition comparisons showed that BHB concentrations increased over the postprandial period only during continuous hypoxemia, reaching 80.5% and 119% higher concentrations at 300 minutes (0.204 mmol/L; 95% CI: 0.175-0.232; P_Bonferroni_ = 0.002) and 360 minutes (0.247 mmol/L; 95% CI: 0.218-0.275; P_Bonferroni_ < 0.001), respectively, than at 60 minutes (0.113 mmol/L; 95% CI: 0.084-0.142). No within-condition differences were observed at any other time points across all conditions (all P_Bonferroni_ ≥ 0.234). Between-condition comparisons at each time point indicated that BHB concentrations at 360 minutes were 13.0% and 14.2% higher during continuous hypoxemia compared with the corresponding time point under normoxemia (0.176 mmol/L; 95% CI: 0.153-0.200; P_Bonferroni_ = 0.029) and intermittent hypoxemia (0.163 mmol/L; 95% CI: 0.139-0.186; P_Bonferroni_ = 0.002), respectively. No between-condition differences were observed at earlier postprandial time points (all P_Bonferroni_ ≥ 0.135).

**Figure 1.**
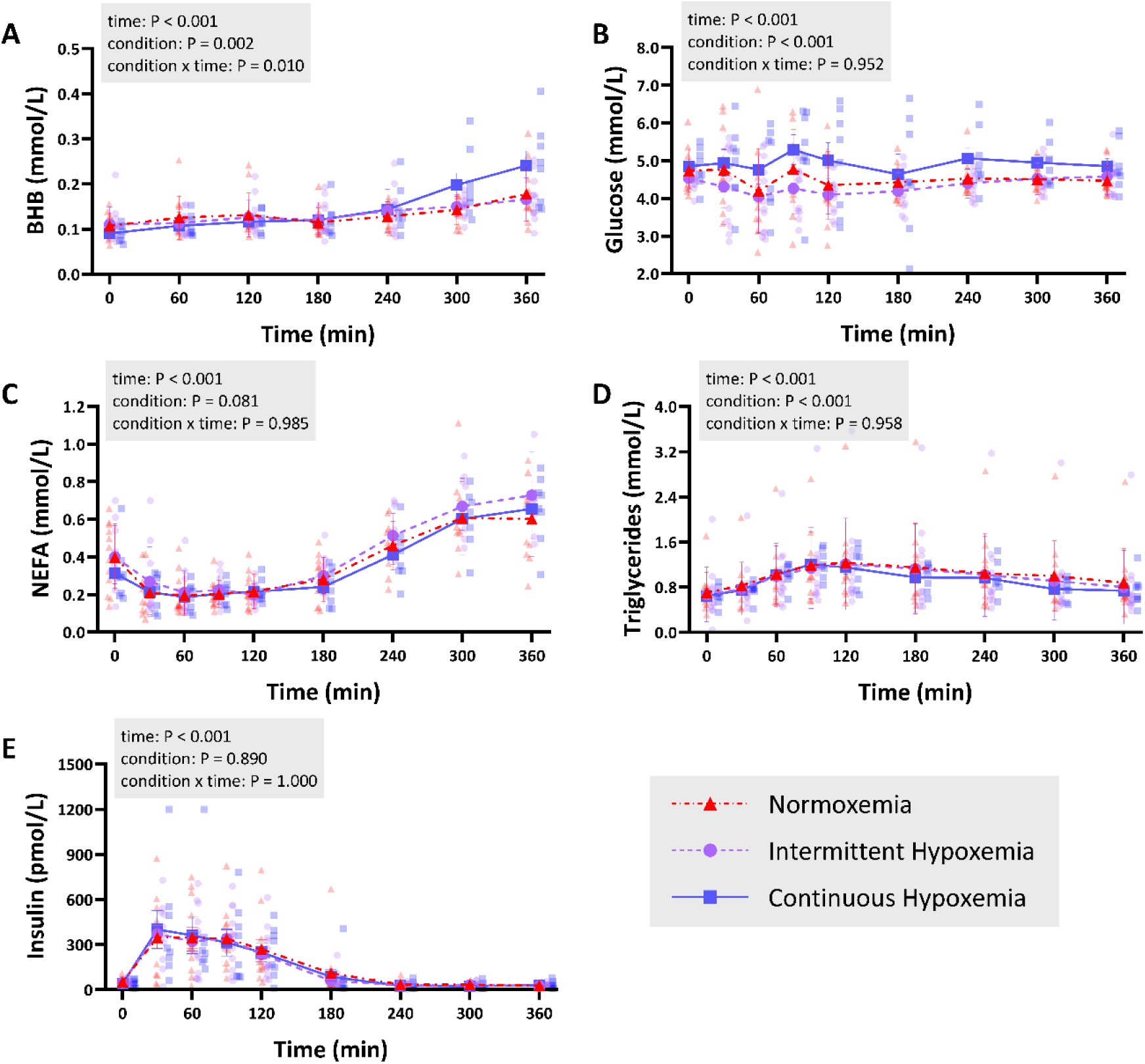
Plasma concentrations of BHB (A), glucose (B), NEFA (C), triglycerides (D), and insulin (E) in young adult females at baseline (0 minutes) and every hour after consuming a high-fat meal during six hours of exposure to normoxemia (red), intermittent hypoxemia (purple), and continuous hypoxemia (blue). Data are presented as means (standard deviations) of raw data and individual points. Postprandial data were analyzed using linear mixed-effects models with condition and time as fixed effects, participant identification as a random intercept, and baseline as a continuous covariate. Abbreviations: β-hydroxybutyrate (BHB), non-esterified fatty acid (NEFA). Note: Model-derived estimated marginal means with 95% confidence intervals are presented in Figure S1 for visualization of baseline-adjusted condition-level effects.

A time × condition interaction (P = 0.003) was also observed when comparing BHB concentrations at baseline and 360 minutes post-meal. Post hoc analyses showed that BHB concentrations were 66.4%, 50.9%, and 172% higher after 360 minutes than at baseline during normoxemia (0 minutes vs 360 minutes: 0.107 mmol/L; 95% CI: 0.078-0.137; vs 0.178 mmol/L; 95% CI: 0.149-0.207; P_Bonferroni_ = 0.003), intermittent hypoxemia (0.110 mmol/L; 95% CI: 0.081-0.140; vs 0.166 mmol/L; 95% CI: 0.136-0.195; P_Bonferroni_ = 0.042), and continuous hypoxemia (0.088 mmol/L; 95% CI: 0.053-0.123; vs 0.239 mmol/L; 95% CI: 0.204-0.273; P_Bonferroni_ < 0.001), respectively.

### 3.3. Plasma metabolic and insulinemic responses to a high-fat meal during six hours of exposure to normoxemia, intermittent hypoxemia, and continuous hypoxemia

Figure 1B-E presents baseline and postprandial plasma concentrations of glucose, NEFA, triglycerides, and insulin. Baseline plasma metabolite and insulin concentrations were similar across experimental conditions (all P ≥ 0.088). For plasma glucose concentrations (Figure 1B), no main effect of time (P = 0.231) or time × condition interaction (P = 0.952) was observed. A main effect of condition was detected (P < 0.001), and post hoc comparisons revealed that mean postprandial glucose concentrations were higher by 9.8% and 12.5% during continuous hypoxemia (4.92 mmol/L; 95% CI: 4.65-5.18) compared to normoxemia (4.48 mmol/L; 95% CI: 4.25-4.71; P_Bonferroni_ = 0.001) and intermittent hypoxemia (4.37 mmol/L; 95% CI: 4.14-4.60; P_Bonferroni_ < 0.001), respectively, whereas normoxemia and intermittent hypoxemia were similar (P_Bonferroni_ = 0.945). For plasma NEFA concentrations (Figure 1C), a main effect of time was observed (P < 0.001), characterized by an early postprandial suppression followed by a progressive rebound later in the postprandial period. There was no main effect of condition (P = 0.081) nor time × condition interaction (P = 0.985), indicating similar NEFA responses across experimental conditions.

For plasma triglyceride concentrations (Figure 1D), a main effect of time was observed (P < 0.001), reflecting the postprandial rise and subsequent decline following the high-fat meal. A main effect of condition was also detected (P < 0.001); however, there was no time × condition interaction (P = 0.958). Post hoc pairwise comparisons showed that, when averaged across the postprandial period, a 14.1% and 13.9% increase in triglyceride concentrations was observed during continuous hypoxemia (1.095 mmol/L; 95% CI: 0.996-1.190) compared to normoxemia (0.960 mmol/L; 95% CI: 0.870-1.050; P_Bonferroni_ = 0.002) and intermittent hypoxemia (0.961 mmol/L; 95% CI: 0.871-1.050; P_Bonferroni_ = 0.001), respectively, with no difference between normoxemia and intermittent hypoxemia (P_Bonferroni_ = 1.000). For plasma insulin concentrations (Figure 1E), a main effect of time (P < 0.001) was observed, whereas no main effect of condition (P = 0.890) or time × condition interaction (P = 1.000) was observed.

## 4. Discussion

This study evaluated the effects of normoxemia, intermittent hypoxemia, and continuous hypoxemia on postprandial plasma BHB concentrations and related cardiometabolic markers in young adult females following a high-fat meal. The primary and novel finding is that continuous hypoxemia, but not intermittent hypoxemia, resulted in higher postprandial BHB concentrations than normoxemia, particularly during the late postprandial period (six hours post-meal). This elevation occurred alongside higher postprandial glucose concentrations and marginally, but statistically, higher postprandial triglyceride concentrations, whereas NEFA and insulin concentrations remained similar across experimental conditions. Collectively, these findings suggest that sustained reductions in oxygen availability acutely alter postprandial metabolism in young adult females.

### 4.1. The effect of normoxemia, intermittent hypoxemia, and continuous hypoxemia on postprandial plasma BHB concentrations after a high-fat meal

Our observation that a high-fat meal increases plasma BHB concentrations above baseline under normoxemia is consistent with previous reports. For example, Halkes et al. (2003) and Meijssen et al. (2000) demonstrated that an oral fat-loading test increased BHB concentrations, with greater increases observed in individuals with familial combined hyperlipidemia, a disorder characterized by impaired postprandial triglyceride and NEFA clearance. These findings suggest that greater hepatic NEFA delivery during the postprandial period promotes hepatic ketone body production. Notably, these increases occur despite the concomitant rise in insulin and suppression of circulating NEFA, which may inhibit ketogenesis (Cotter et al., 2013). Whether postprandial changes in BHB clearance also contribute to this response remains unclear.

The present study extends these observations by demonstrating that six hours of continuous hypoxemia increases late postprandial plasma BHB concentrations in young adult females after a high-fat meal, compared with normoxemia and intermittent hypoxemia, despite similar circulating NEFA and insulin responses across conditions. This design allowed a direct comparison of hypoxemia patterns under matched postprandial conditions, which was not possible in our previous work (Marcoux et al., 2022). In that study, continuous hypoxemia increased BHB concentrations in young adult males during a six-hour fast, whereas intermittent hypoxemia in the postprandial state had no statistically significant effect compared with normoxemia (Marcoux et al., 2022). However, differences in nutritional status precluded direct comparisons of hypoxemia patterns, and the statistical approach used a repeated-measures ANOVA rather than a linear mixed-effects model. The concomitant elevations in postprandial plasma triglyceride and glucose concentrations during continuous hypoxemia may help explain the BHB response observed in the present study. Elevated postprandial triglyceride concentrations may reflect impaired systemic triglyceride clearance or increased triglyceride production, both of which may indicate greater hepatic NEFA availability, which could lead to increased hepatic BHB production (Cotter et al., 2013). In parallel, elevated postprandial glucose concentrations may reflect subtle alterations in hepatic glucose uptake, storage, or production, or in insulin sensitivity, that, although insufficient to alter circulating insulin concentrations, could influence hepatic substrate partitioning (Newhouse et al., 2017; Woolcott et al., 2015). Together, these findings suggest that continuous hypoxemia alters postprandial BHB concentrations through mechanisms not fully explained by circulating NEFA or insulin concentrations alone.

In contrast to continuous hypoxemia, exposure to six hours of intermittent hypoxemia did not result in higher postprandial plasma BHB concentrations relative to normoxemia. The lack of effect under intermittent hypoxemia may be partly explained by inherent differences in hypoxemic dose (i.e., different levels of oxygen deprivation) or by the distinct molecular mechanisms, such as the regulation of hypoxia-inducible factor, that are unique to intermittent hypoxemia as opposed to those regulated by continuous hypoxemia (Hunyor & Cook, 2018). However, these explanations remain speculative and further research is warranted. Additionally, sex-specific metabolic responses may further modulate postprandial BHB responses. For example, previous findings show that females have higher postprandial BHB concentrations than males, potentially due to greater postprandial hepatic NEFA oxidation in females (Halkes et al. 2003).

### 4.2. Limitations

Several limitations should be acknowledged. First, this study represents a secondary analysis of a larger randomized crossover trial, and an a priori power calculation was not performed for BHB responses. As such, these findings should be interpreted as exploratory. Second, the continuous hypoxemia condition was conducted as part of a follow-up session and was therefore not randomized nor restricted to a single menstrual cycle phase. Third, participants were not blinded to the experimental conditions. Finally, the findings are specific to the controlled laboratory conditions employed, including passive rest and simulated hypoxemic exposures, and may not fully generalize to more ecologically valid settings such as active high-altitude ascent or habitual sleep-related hypoxemia in free-living environments.

### 4.3. Conclusion

In conclusion, this study demonstrates that continuous hypoxemia, but not intermittent hypoxemia, increases postprandial plasma BHB concentrations in young adult females following a high-fat meal, relative to normoxemia. This effect emerged primarily during the late postprandial period and occurred independently of changes in plasma NEFA and insulin concentrations, two key regulators of BHB production. More broadly, these findings highlight the importance of considering the characteristics of hypoxemic exposure, including its pattern and dose, when evaluating postprandial metabolic responses under hypoxemia. Future studies are warranted to investigate how hypoxemia may alter circulating ketone bodies and to determine whether elevated BHB concentrations confer metabolic advantages when oxygen supply is limited in humans.

## Data Availability Statement

The datasets generated for this study are available upon reasonable request to the corresponding author and with a signed access agreement.

## Author Contributions

All authors approved the final version of the manuscript; agree to be accountable for all aspects of the work in ensuring that questions related to the accuracy or integrity of any part of the work are appropriately investigated and resolved; and all persons designated as authors qualify for authorship, and all those who qualify for authorship are listed. **Nicholas Goulet:** Methodology, Formal Analysis, Investigation, Writing – Original Draft, Writing – Review & Editing, Visualization. **Alexanne Laroque:** Formal Analysis, Investigation, Writing – Original Draft, Writing – Review & Editing. **Caroline Marcoux:** Investigation, Writing – Review & Editing. **Vincent Bourgon:** Investigation, Writing – Review & Editing. **Jean-François Mauger:** Methodology, Investigation, Writing – Review & Editing. **Ruwan Amaratunga:** Funding Acquisition, Writing – Review & Editing. **Pascal Imbeault:** Conceptualization, Methodology, Writing – Review & Editing.

## Acknowledgements

We thank all the individuals who participated in this study and acknowledge the support of the Behavioural and Metabolic Research Unit members who helped with data collection. We thank the Registered Nurses at the University of Ottawa for their assistance.

All authors have read and approved this version of the manuscript. This article was last modified on March 11, 2026.

