## Supplemental Figure 1 for "Plasma β-hydroxybutyrate Concentrations in Young Adult Females After a High-Fat Meal Under Normoxemia, Intermittent Hypoxemia, and Continuous Hypoxemia"

<u>**Supplementary File**</u>

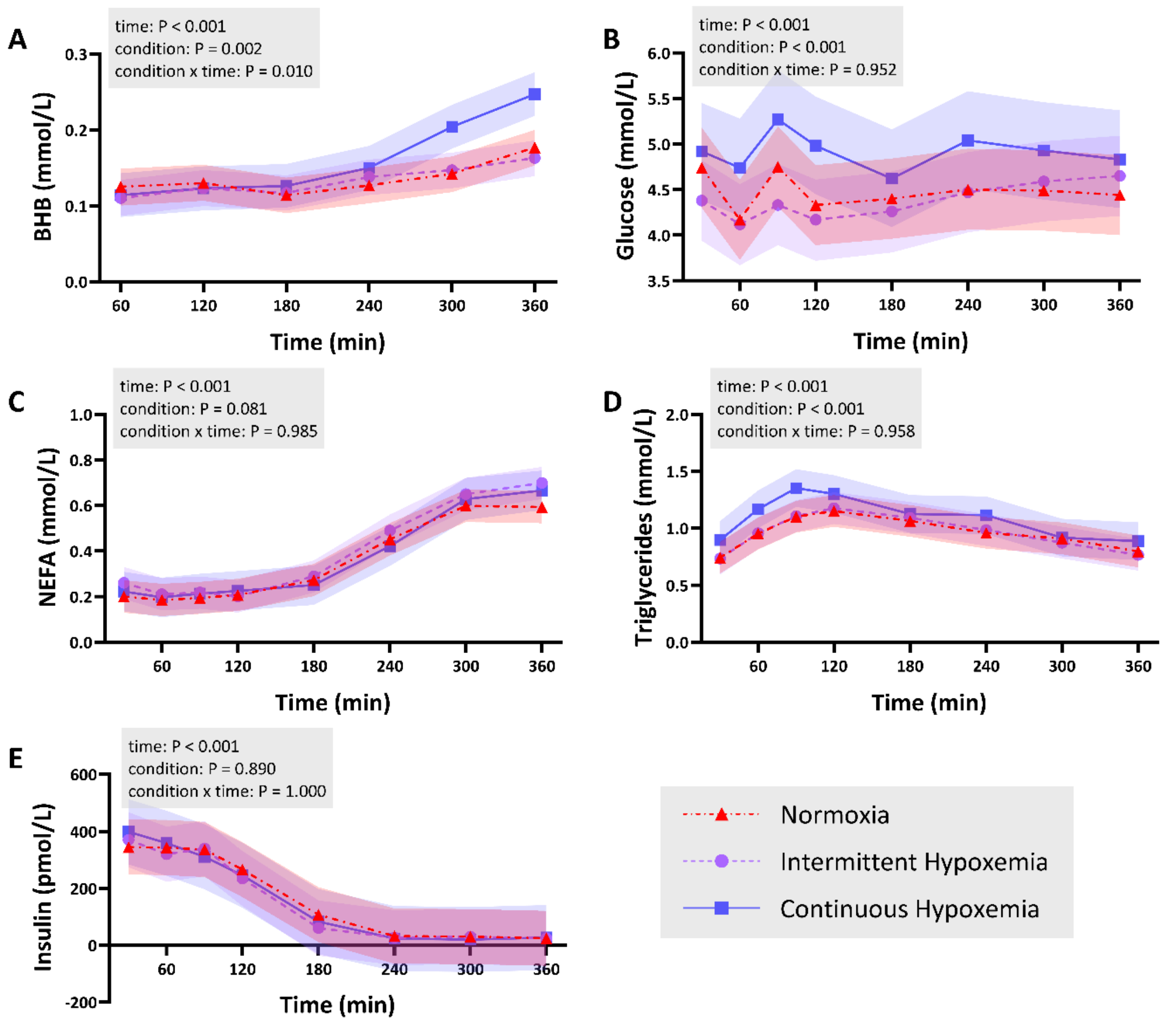


**Figure S1.** Postprandial plasma concentrations of BHB (A), glucose (B), NEFA (C), triglycerides (D), and insulin (E) in young adult females after consuming a high-fat meal during six hours of exposure to normoxemia (red), intermittent hypoxemia (purple), and continuous hypoxemia (blue). Data are presented as model-derived estimated marginal means (95% confidence intervals). Postprandial data were analyzed using linear mixed-effects models with condition and time as fixed effects, participant identification as a random intercept, and baseline (0 minutes) as a continuous covariate. Abbreviations: β-hydroxybutyrate (BHB), non-esterified fatty acid (NEFA).
